# Cumulative multisensory discrepancies shape the ventriloquism aftereffect but not the ventriloquism bias

**DOI:** 10.1101/2022.09.06.506717

**Authors:** Christoph Kayser, Hame Park, Herbert Heuer

## Abstract

Multisensory integration and recalibration are two processes by which perception deals with discrepant signals. Both are often studied in the spatial ventriloquism paradigm. There, integration is probed by the presentation of discrepant audio-visual stimuli, while recalibration manifests as an aftereffect in subsequent unisensory judgements. Both biases are typically quantified against the degree of audio-visual discrepancy, reflecting the possibility that both may arise from common underlying multisensory principles. We tested a specific prediction of this: that both processes should also scale similarly with the history of multisensory discrepancies experienced in previous trials. Analysing data from ten experiments we confirmed the expected dependency of each bias on the immediately presented discrepancy. And in line with the aftereffect being a cumulative process, this scaled with the discrepancies presented in multiple preceding audio-visual trials. However, the ventriloquism bias did not depend on the history of multisensory discrepancies and also did not depend on the aftereffect biases in previous trials - making these two multisensory processes experimentally dissociable. These findings support the notion that the ventriloquism bias and the aftereffect reflect distinct functions, with integration maintaining a stable percept by reducing immediate sensory discrepancies and recalibration maintaining an accurate percept by accounting for consistent discrepancies.

## Introduction

Our brain combines multisensory signals to guide immediate behaviour, but discrepant multisensory signals can also exert lasting influences even on subsequent unisensory judgments. A prototypical example is the spatial ventriloquism paradigm: here the discrepant positions of visual and acoustic stimuli are combined when localizing a sound source - the ventriloquism effect. In addition, both signals can influence the localization of subsequent unisensory acoustic stimuli – the ventriloquism aftereffect (Recanzone, 2009; Wozny and Shams, 2011b; Frissen et al., 2012; Bruns and Roder, 2015; Mendonça et al., 2015; Bosen et al., 2017; Bosen et al., 2018; Park and Kayser, 2019; Badde et al., 2020). This aftereffect – or recalibration bias - emerges in the absence of explicit task feedback and on multiple time scales. Importantly, both integration and recalibration are typically described by their dependency on the spatial discrepancy presented in the multisensory trials. Their similar dependency on this multisensory dimension can be taken to suggest that both arise from a common underlying multisensory mechanism.

Indeed, one line of work supports the notion that the aftereffect is a direct consequence of the preceding integration of multisensory signal (Bruns, 2019; Rohlf et al., 2020; Noppeney, 2021). For example, the discrepancy between integrated multisensory signals and subsequent unisensory stimuli apparently drives recalibration (Wozny and Shams, 2011b; Bruns and Roder, 2015; Park and Kayser, 2019; Rohlf et al., 2020; Noppeney, 2021), and both biases are strongest when the multisensory stimuli are judged as being causally related (Wozny and Shams, 2011b; Park and Kayser, 2020). Furthermore, both are similarly affected by manipulations of stimulus reliability (Rohlf et al., 2021) and attention (Badde et al., 2020), and neuroimaging studies have pointed to partly overlapping neurophysiological processes that shape integration and recalibration (Park and Kayser, 2019, 2021). If this notion were correct, experimental manipulations affecting integration should also affect recalibration. For example, both biases should depend in a similar manner on the history of the multisensory experience, such as the discrepancies between the acoustic and visual signals.

An alternative view stipulates that recalibration is independent of whether the preceding multisensory signals are deemed causally related but is rather shaped by the believe in a modality-specific bias (Di Luca et al., 2009; Block and Bastian, 2011; Burr et al., 2011; Wozny and Shams, 2011a; Zaidel et al., 2013; Bosen et al., 2017; Noppeney, 2021). Still, this believe in a modality specific bias may be driven by multisensory evidence, hence the scaling of the aftereffect with the degree of multisensory discrepancy. If this view were correct, integration and recalibration could be differentially affected by the history of multisensory stimuli. While some previous studies have probed the history dependence of the aftereffect, these either did not provide a direct comparison of integration and recalibration biases or featured experimental designs that were limited by the use of a fixed and hence predictable spatial discrepancy (Wozny and Shams, 2011b; Mendonça et al., 2015; Bosen et al., 2017).

To arbitrate between these predictions, we analysed data from ten experiments and directly tested whether the spatial ventriloquism effect and the aftereffect depend in a similar manner on the history of multisensory discrepancies experienced. All experiments featured a prototypical series of audiovisual and auditory trials (AV-A) that in the original experiments was used to probe the scaling of integration (in the AV trial) and of immediate recalibration (A trial) with the spatial discrepancy presented in the immediate AV trial. Importantly, the direction and magnitude of the multisensory spatial discrepancies was variable and unpredictable from trial to trial. We here leveraged this data to model both biases against the history of multisensory discrepancies across multiple previous AV trials. Our results provide converging and strong evidence that the aftereffect scales in a cumulative fashion with the series of experienced discrepancies, while multisensory integration in the form of the ventriloquism bias does not. This confirms the cumulative nature of the aftereffect and shows that the ventriloquism bias is tied to the immediately present but not previous degrees of multisensory discrepancies.

## Methods

### Experimental design and procedures

We analysed data from 10 experiments, eight of which have been published previously (see Table 1 for details). All experiments followed the same overall design template to probe the ventriloquism effect and the immediate aftereffect following established work (Wozny and Shams, 2011b; Park and Kayser, 2019, 2020, 2021; Park et al., 2021; Park and Kayser, 2022). Participants were adult volunteers who reported no history of neurological diseases, and normal vision and hearing. They were compensated for their time and provided written or electronic informed consent prior to participation. Before the actual experiments we performed a screening task to probe participants’ spatial hearing thresholds (Park and Kayser, 2022). The studies were approved by the local ethics committee of Bielefeld University and data were collected anonymously. This makes it impossible to determine whether some participants participated in more than one of the experiments reported here.

**Table 1:** Datasets. The datasets analysed here come from 10 experiments that capitalize on the same overall spatial ventriloquism design. Experiments 1-8 have been published as part of previous studies. Experiments 9 and 10 are analysed for the first time and contained additional manipulations not analysed here. N indicates the number of participants. The last three columns indicate the spatial separation of the five stimulus locations, the number of AV-A trial-pairs per participant and the fraction of V only trials that were not analysed here.

| Experiment | Publication & DOI | Note | N participants | Stimulus spacing | AV-A pairs | V trials |
| --- | --- | --- | --- | --- | --- | --- |
| 1 | Park & Kayser. The Journal of Neuroscience 2021, doi: 10.1523/JNEUROSCI.2091-20.2020 | Short-term recalibration data only | 19 | 11.6° | 360 | 7% |
| 2 | Park, Kayser. Scientific Reports 2020, doi: 10.1038/s41598-020-77730-7 | Experiment 1 | 20 | 8° | 375 | 10% |
| 3 | Park & Kayser C. Scientific Reports 2020, doi: 10.1038/s41598-020-77730-7. | Experiment 2 | 21 | 8° | 375 | 10% |
| 4 | Park & Kayser C. Scientific Reports 2020, doi: 10.1038/s41598-020-77730-7. | Experiment 3 - mask | 22 | 8° | 180 | 10% |
| 5 | Park & Kayser C. Scientific Reports 2020, doi: 10.1038/s41598-020-77730-7. | Experiment 3 – no mask | 22 | 8° | 180 | 10% |
| 6 | Park, Nannt & Kayser. Cortex 2021, doi: 10.1016/j.cortex.2020.12.001 | Young participants only | 22 | 8.5° | 396 | 8% |
| 7 | Park & Kayser C. Cognition 2022, doi: j.cognition.2022.105092 | Narrow context | 21 | 6.7° | 396 | 8% |
| 8 | Park & Kayser C. Cognition 2022, doi: j.cognition.2022.105092 | Wide context | 21 | 11.6° | 396 | 8% |
| 9 | Previously unpublished | Trial context manipulation | 17 | 11° | 465 | 12% |
| 10 | Previously unpublished | Naturalistic stimuli | 20 | 11° | 360 | 15% |

In all experiments, the auditory and visual stimuli were presented either simultaneously in multisensory trials (AV) or individually in unisensory trials (A or V; c.f. Figure 1A). The experiments were performed in a dark and anechoic chamber with sounds being presented at 64 dB r.m.s. through one of 5 speakers (Monacor MKS-26/SW, MONACOR International GmbH & Co. KG, Bremen, Germany) located at 5 horizontal locations. The precise spacing of stimulus locations differed between experiments (Table 1). The visual stimuli were projected (Acer Predator Z650, Acer Inc., New Taipei City, Taiwan) onto an acoustically transparent screen placed in front of the speakers (Screen International Modigliani, 2×1 m^2^) and were centred on the same five locations as the sounds. The positions of auditory and visual stimuli were drawn pseudo randomly to sample a large range of multisensory discrepancies (the spatial separation of the auditory and visual stimuli in the AV trials) and stimulus positions. This made the direction of the spatial discrepancy presented in each trial and the magnitude of this unpredictable to the participants, unlike in some other previous studies that used a fixed magnitude or a fixed direction of spatial discrepancy for each participant (Mendonça et al., 2015; Bosen et al., 2017). The participants were asked to localize the sound (on AV and A trials) or to localize the visual stimulus (on V trials). They responded by moving a mouse cursor to the respective location and clicking to submit their response. The AV trials served to probe the ventriloquism bias and to induce the aftereffect; the A trials served to probe the aftereffect; the less frequent V trials served to ensure that participants maintained divided attention across both senses. The AV trials were always followed by A trials, which were sometimes followed by V trials. The overall number of trials and the proportion of V trials differed between experiments (see Table 1).

**Figure 1:**
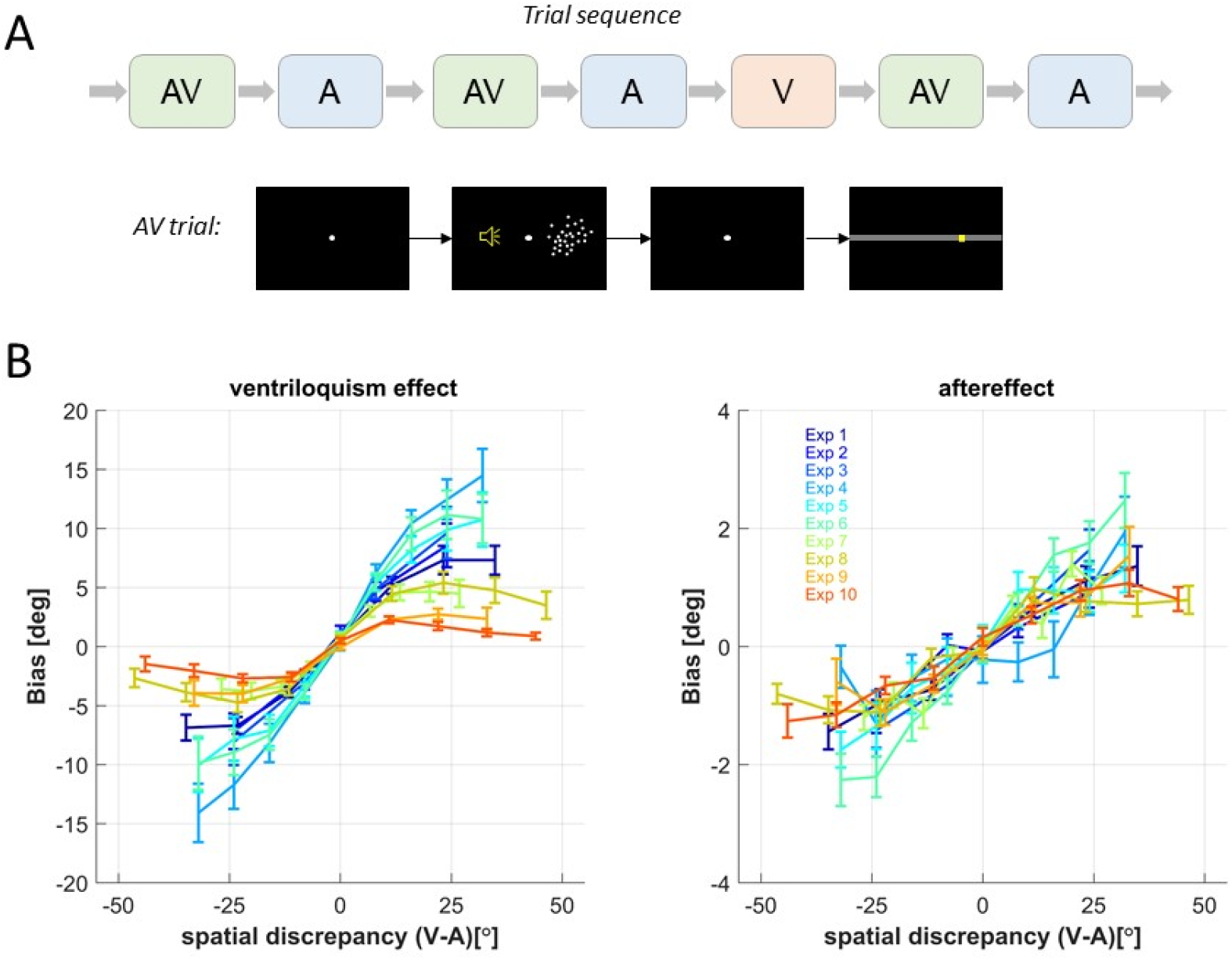
Experimental design and response biases. **A)** shows the typical sequence of AV and A trials designed to probe the two responses biases, and the less frequent V trials to maintain attention on both modalities. The lower panel in A shows a typical sequence of events within an AV trial, including a fixation period, stimulus presentation (here random dots), a post-stimulus period and the response bar on which participants moved a mouse cursor to indicate the perceived location of auditory or visual stimuli. The yellow speaker icon indicates the location of a sound, but was not displayed in the experiment. **B)** shows the average (mean and s.e.m.) bias for each of ten experiments as a function of the spatial discrepancy presented in the AV trial probing the ventriloquism effect or the AV trial immediately prior to the A trial probing the aftereffect.

#### Stimuli experiments 1-9

The visual stimulus was a cloud of white dots distributed according to a twodimensional Gaussian distribution (200 dots, SD of vertical and horizontal spread: 1.6° (Experiments 2-5,7,8) or 2° (Experiments 1,6,9), width of a single dot: 0.12°, duration of stimulus presentation: 50 ms), and the auditory stimulus was a 1300 Hz sine wave tone (50 ms duration). Experiment 9 was originally designed to probe the impact of trial-context and presented the AV, A and V trials in three sub-blocks featuring different proportions of AV trials with a visual stimulus being displaced from the sound towards the center of the visual screen. For the present analysis, however, we combined all trials and ignored this specific manipulation.

#### Stimuli experiment 10

For this experiment we used naturalistic images and sounds as stimuli. More precisely, the visual stimuli were either the faces of one of five animals (cat, dog, eagle, sheep, tiger) or phase-scrambled versions of these images. The images were faded into the grey background using a Gaussian mask with a horizontal SD of 2° and were equalized for luminance and contrast. The acoustic stimuli were the matching five vocalizations (with 200 ms duration, all r.m.s normalized) or scrambled versions of these. These scrambled sounds were created by filtering the original sounds into 10 equi-spaced band-limited components between 100Hz and 8kHz and randomly interchanging these bands across sounds. AV trials either presented matching naturalistic auditory and visual pairs, scrambled pairs, or pairs featuring one naturalistic and one scrambled stimulus (again these specific manipulations are neglected for the present analyses).

### Analysis of response biases

We used identical definitions of the trial-wise ventriloquism bias and the aftereffect to facilitate a direct comparison of these. Each bias was defined as the difference between the reported sound location (in the AV trial for the ventriloquism bias, the A trial for the aftereffect) and the average reported location in all A trials for this specific sound position. This definition ensures that other response tendencies, such as to report stimuli closer to the midline as they are (also known as central bias), or to report them slightly shifted towards either side, are equally accounted for in the ventriloquism effect and the aftereffect (Wozny and Shams, 2011b; Mendonça et al., 2015; Rohe and Noppeney, 2015b; Park et al., 2021). Typically, the trial-wise biases are then investigated in relation to the multisensory discrepancy in the respective AV trial in which the ventriloquism effect is obtained, or the AV trial immediately prior to the A trial in which the ventriloquism aftereffect is obtained (Figure 1B). We here extended this analysis and probed each bias in relation to the discrepancies in up to three successive AV-A trial-pairs. For this analysis we ignored the fact that some AV-A trial-pairs were interspersed with visual only trials, given that these visual stimuli should not affect auditory spatial recalibration and given that the ventriloquism bias has been shown to be robust against intervening mnemonic manipulations or other additional unisensory judgements (Bosen et al., 2017; Park and Kayser, 2020).

Both biases scale with the sign and magnitude of the spatial discrepancy (Figure 1B for experimentwise averages). This dependency can taper off at large discrepancies, reflecting the reduced combination of spatially very discrepant signals. Such a non-linear dependency is predicted by models of multisensory causal inference, which posit that in a typical ventriloquism setting multisensory signals are only combined when they are likely to arise from a common source (Kording et al., 2007; Odegaard et al., 2015; Rohe and Noppeney, 2015a; Noppeney, 2021; Shams and Beierholm, 2022). To account for such nonlinear dependencies, we included both linear and nonlinear predictors in the analyses described below, analogously to our previous work (Park and Kayser, 2020, 2021; Park et al., 2021; Park and Kayser, 2022): for a given spatial discrepancy, Δva, the non-linear term was modelled as sign(Δva) * sqrt(Δva).

We quantified the dependency of each bias on the spatial discrepancies in up to three AV-A trial-pairs, as follows. In a first analysis, we compared proportional biases contingent on different constellations of the sign of the spatial discrepancy in preceding trial-pairs (Figure 2A). For this we focused on sequences of trial-pairs (denoted by a,b,c in the following) that all featured a non-zero spatial discrepancy. The proportional bias was computed as the ratio of the trial-wise bias observed in pair c relative to the discrepancy in trial-pair c. We derived this for the following constellations of trials: across all trial-pairs (‘c’ in Figure 2B), for sequences of two pairs with same (‘b=c’) or opposing (‘b~=c’) signs of the discrepancy, and for triplets with the same sign (‘a=b=c’) or a change in sign (‘a=b~=c’). We implemented this analysis as it allows a direct visualization of the dependence of each bias on the history of spatial discrepancies.

**Figure 2:**
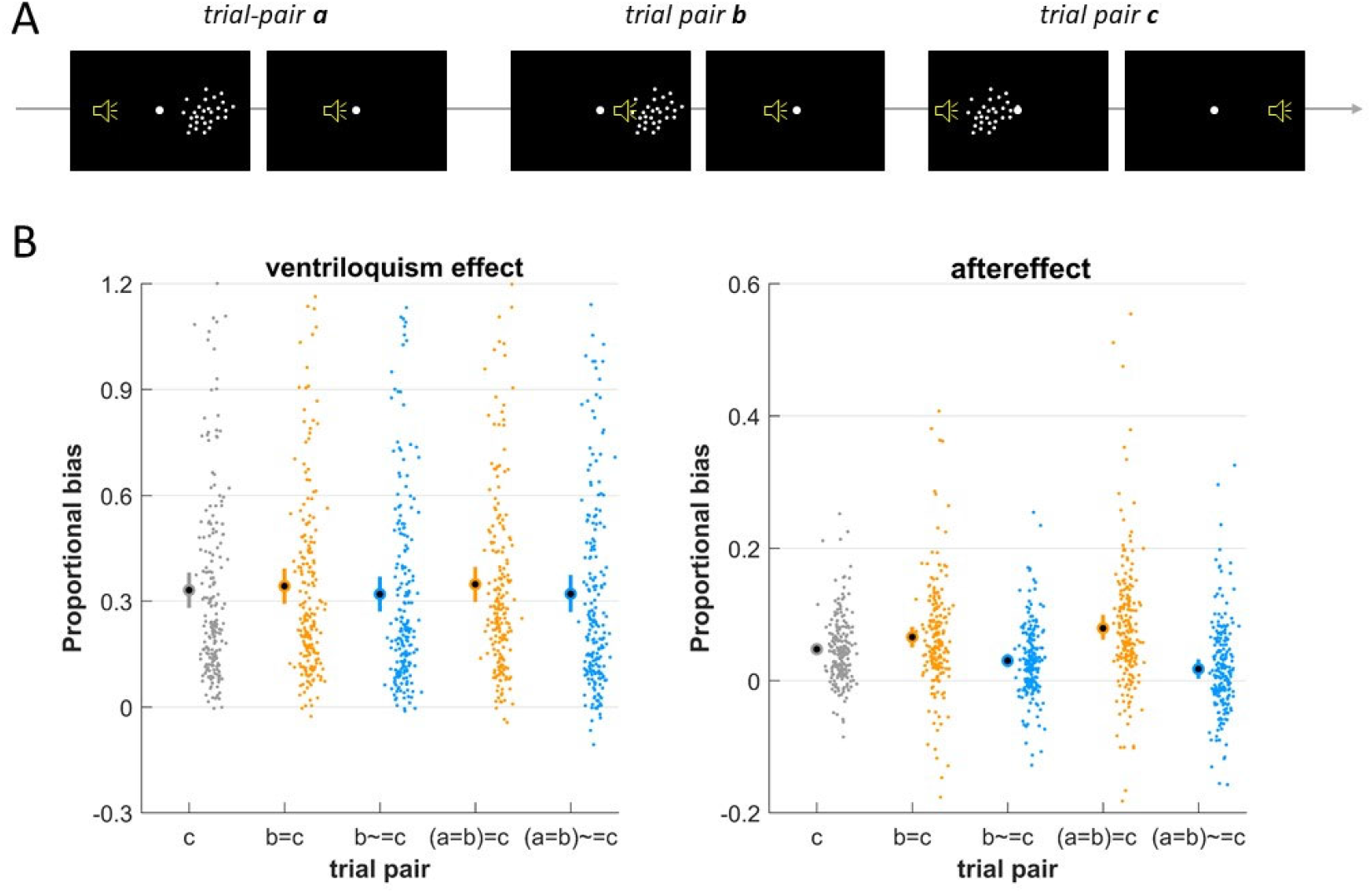
History-dependence is prominent for the aftereffect but not for the ventriloquism effect. **A)** For analysis we probed the dependency of each bias obtained in a given AV-A trial-pair (termed c) on the spatial discrepancy in that trial-pair and two preceding trial pairs (here labelled a and b). For the analysis in this figure we grouped trial-pairs based on the direction of the spatial discrepancy, for analyses reported below we included also the precise numeric values. **B)** Proportional biases contingent on the history of the sign of the multisensory discrepancy. The bias was quantified for individual AV-A pairs (condition c), sequences of two pairs with same (‘b=c’) or opposing sign of discrepancy (‘b~=c’), triplets featuring the same sign (‘a=b=c’) and triplets featuring a change in sign (‘a=b~=c’). The criterion of an equal sign of the discrepancy selects trials on which the visual stimulus is always on the same side of the auditory stimulus, regardless of their absolute locations. The proportional bias was defined as the ratio of trial-wise bias relative to the discrepancy in trial-pair c. Dots reflect individual datasets (i.e. participants, n=205), circles the overall mean and error bars the 99% bootstrap confidence interval.

In a second analysis, we fit participant-wise linear models to predict each trial-wise bias (in trial-pair c) based on the precise discrepancies in this immediate and previous trial-pairs. These models included both linear and non-linear predictors for the discrepancies and included also trials with zero discrepancy (in contrast to the above analysis of proportional biases). Specifically, we modelled the trial-wise bias based on the immediate discrepancy (in trial-pair c), by also including one previous pair (b&c) or by including two preceding pairs (a&b&c). We then compared the explained variance (R^2^) between these models across participants. In addition, we used variance partitioning to quantify the variance that is uniquely explained by the immediate discrepancy (c) and the preceding discrepancies (a,b). Variance partitioning splits the explained variance into proportions uniquely explained by individual predictors (also known as semi-partial correlations) and those common to two or more predictors (SEIBOLD and McPHEE, 1979; Freckleton, 2002; Nimon et al., 2008). In the present data these common variances were mostly negligible. Because fitting linear models with multiple predictors on limited data can result in biased estimates of variance, and because individual experiments contained different numbers of trials, we corrected the obtained variance estimates: for each model we obtained an estimate of the statistical bias (i.e. the variances explained by chance) by fitting the models 500 times after randomizing the dependent variable across trials (Benjamini and Yu, 2013). The median of these 500 variance estimates was then subtracted from the original estimates. As a result of this bias correction, the corrected variance estimates can be negative, with small values indicating near chance-level results.

In a third analysis, we entered the combined data across all ten experiments using single linear mixed effect models for each bias. We fit models including only the immediate trial-pair (c) or also including the discrepancies of one or two previous trial-pairs. These models again included both linear and nonlinear predictors for each discrepancy and included random offsets and slopes for the individual discrepancy predictors for each experiment. For each bias we then compared the predictive power of the three models (including trials c, b&c, a&b&c) using their BIC values (Raftery, 1995; Gelman et al., 2014). Given the large number of predictors, this analysis may underestimate the relevance of previous trials as models including three trial-pairs may be overly punished for their high number of predictors. Still, we found that this analysis and that using models fit to individual datasets above yield the same overall conclusions.

### Statistical analysis

Statistical comparisons are based on the 205 datasets analysed, which we treated as independent samples, though it is possible that some participants completed more than one experiment. To describe condition-wise effects across the sample we relied on population means and 99% (percentile) bootstrap confidence intervals obtained using 5000 randomizations (Fethney, 2010). For a statistical assessment of differences between conditions we relied on paired t-tests for which we report Cohens’ D as effect size and Bayes factors (BF) as indicators of significance. Because the present study emphasizes the qualitative and quantitative comparison of the two perceptual biases we refrain from reporting p-values but emphasize effect sizes and Bayesian statistics (Wagenmakers, 2007; Rouder et al., 2009; Greenland et al., 2016). When interpreting BF’s we refer to the nomenclature of Raftery (Raftery, 1995): we interpreted BF between 1 and 3 as ‘weak’, between 3 and 20 as ‘positive’ and between 20 and 150 as ‘strong’ evidence.

## Results

We analysed the ventriloquism effect and the immediate aftereffect in 205 datasets that were collected across ten variations of the same overall experimental design (see Table 1). This design consisted of a sequence of alternating audio-visual (AV) and auditory (A) trials, interspersed with visual trials that served to maintain attention divided across modalities (Figure 1A). In this design the AV trials allow quantifying the ventriloquism effect as a function of the multisensory spatial discrepancy presented in that trial. The subsequent A trials allow quantifying the aftereffect as a function of the discrepancy in the immediately preceding AV trial.

Figure 1B shows the experiment-averaged biases as a function of the spatial discrepancy in the respective AV-A trial-pair. These reveal a general increase with increasing discrepancy but also a tapering off at higher discrepancies that is more prominent for some than other datasets and generally more prominent for the ventriloquism effect. This nonlinear scaling is expected based on models of multisensory causal inference, which predict a stronger bias when two stimuli are judged as originating from a common source and a weaker bias when judged as originating from distinct sources (Kording, 2014; Rohe and Noppeney, 2015b). Given that the spatial separation provides one important cue for a common origin, this gives rise to the nonlinear dependency of the bias on discrepancy.

Our main question was whether each bias depends only on the spatial discrepancy experienced in the immediate AV-A trial-pair, or whether these also depend on the discrepancies presented in preceding AV trials. In a first analysis we quantified the proportional biases for sequences of trial-pairs that were selected for specific constellations of the direction of audio-visual discrepancies (Figure 2). Specifically, we compared the proportional biases for sequences featuring the same sign of discrepancy (e.g. ‘b=c’, ‘a=b=c’), or for sequences featuring a change in sign (e.g. ‘a~=b’, ‘a=b~=c’).

This revealed a cumulative influence of discrepancy on the aftereffect that is only weakly present for the ventriloquism effect. Following sequences of three trial-pairs featuring the same direction of discrepancy the ventriloquism bias was largely comparable to that in sequences featuring a change in sign (‘a=b=c’: 0.348 [0.299 0.401], mean and 99% bootstrap CI, ‘a=b~=c’: 0.320 [0.268 0.376]); in contrast, the aftereffect differed considerably (‘a=b=c’: 0.079 [0.062 0.098], ‘a=b~=c’: 0.018 [0.002 0.032]). A direct comparison provided positive evidence for a difference between constellations for the ventriloquism effect with a small effect size (paired t-test ‘a=b~=c’ vs. ‘a=b=c’: BF=10.2, Cohens’ D=0.22), while for the aftereffect the evidence for a difference was very strong and the effect size moderate (BF=10^7^, Cohens’ D=0.46). While this analysis clearly visualizes the differences between biases it was restricted to trial sequences featuring only non-zero discrepancies and did not account for potential non-linear dependence.

Hence for a more complete analysis we included all AV-A trial pairs and modelled the participant-wise biases based on the specific values of the spatial discrepancies in the immediate trial-pair and the two preceding trial-pairs. These models allowed for potential non-linear dependencies of each bias on the discrepancies. Figure 3 shows two results of this analysis. The upper panels in Figure 3A show the total explained variance for each bias when including only the immediate, one or two preceding trial-pairs. For the ventriloquism bias the explained variance was very similar for the three models, while for the aftereffect the explained variance increased when including previous trial-pairs. Indeed, the relative loss of explained variance when ignoring the two preceding trial-pairs was less than ten percent for the ventriloquism effect (comparing model a&b&c to c only: 7.99% [5.75 10.65], mean and CI; Figure 3 B), while it was more than five times for the aftereffect (43.30% [38.38 48.14]). A direct comparison provided very strong evidence that the explanatory power provided by preceding trial-pairs was larger for the aftereffect (paired t-test: Cohens’ D=1.66, BF=10^40^).

**Figure 3:**
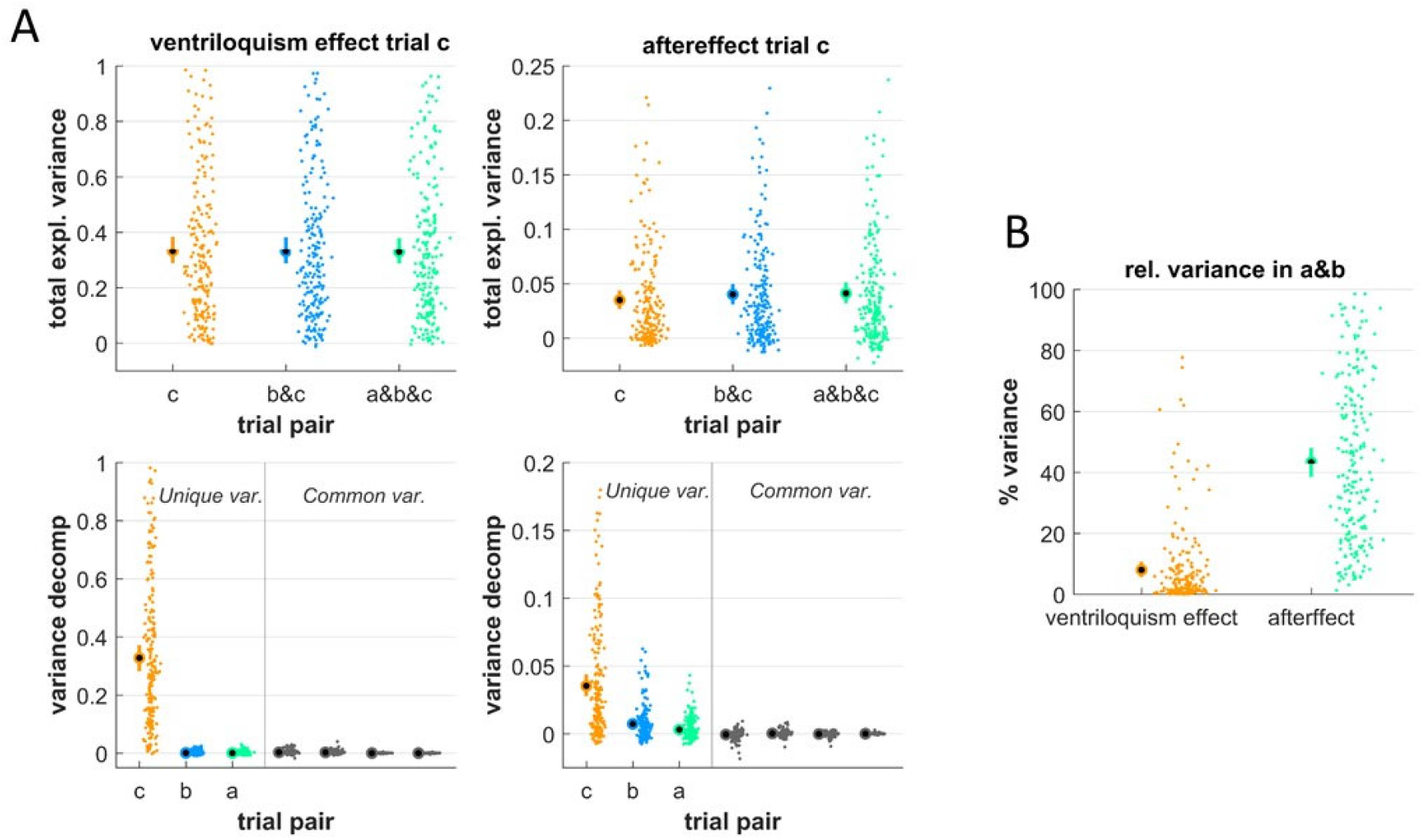
Modelling each bias against the spatial discrepancies in the immediate and preceding trial pairs. For each dataset we modelled the biases against the spatial discrepancy in the immediate AV trial-pair (here labelled c) and one or two preceding trial pairs (a,b). These models featured both linear and non-linear dependencies on each discrepancy to account for the nonlinearities visible in Figure 1. **A)** The upper panels display the total explained variance, the lower panels the results from a variance partitioning analysis. The latter assigns the explained variance to components uniquely explained by individual trials (in colour) or shared variance that is jointly explained by different trials. Here the common variances were all negligible with mean values close to zero and/or 99% CIs reaching zero (from left to right: c and b, c and a, b and a, all three). **B)** Proportion of variance lost when ignoring the preceding trials. This was obtained for each dataset as the different between the model relying only on trial c and that relying on trials a&b&c, expressed relative to the variance for trial c in units of percent. Dots reflect individual datasets (i.e. participants, n=205), circles the overall mean and error bars the 99% bootstrap confidence interval.

In addition, we partitioned the explained variance into contributions uniquely explained by individual trial-pairs and the variance shared by these (Figure 3A, lower panels). The shared contributions were negligible, as expected given the largely independent values of spatial discrepancies across trials in this experimental design. For both biases the unique contribution of the immediate discrepancy was the strongest predictor (ventriloquism effect: 0.328 [0.283 0.374]; aftereffect: 0.035 [0.027 0.044]). Importantly, the contributions of the preceding trials differed between biases: these were negligible for the ventriloquism effect (trial-pair b: 0.001 [0 0.002]; trial-pair c: 0.000 [-0.001 0.001]) but not for the aftereffect (trial-pair b: 0.007 [0.005 0.010]; trial-pair b: 0.003 [0.002 0.005]), although the overall explained variance was larger for the ventriloquism effect. A direct comparison provided very strong evidence that the unique variance explained by the preceding trial-pairs was larger for the aftereffect (trial-pair b: Cohens’ D=0.64, BF=10^7^, trial c: D=0.39, BF=162). Hence, only for the aftereffect did the spatial discrepancy in preceding trial-pairs contribute significant explained variance.

In a third analysis we used linear mixed effects models to predict each bias across the entire sample of datasets based on linear and nonlinear dependencies on the discrepancies across up to three trial-pairs. Again we compared the predictive power of a model including only the immediate trial-pair (c) with models including one (b&c) or two preceding trial-pairs (a&b&c). For the ventriloquism effect the data revealed very strong evidence that they were best explained by a model not including preceding trial-pairs (relative BIC: 0, 26.1 and 105.9), while for the aftereffect we found very strong evidence that the inclusion of a preceding trial-pair provided a better account (relative BIC: 77.5, 0 and 35.8). This corroborates the above results that the ventriloquism effect is tied only to the immediate multisensory discrepancy while the aftereffect is shaped also by those in preceding trials.

### No contribution of the aftereffect to subsequent integration

The first analysis above shows that numerically both response biases may increase following a series of trials with a common direction of spatial discrepancy, though the increase was much more pronounced for the aftereffect. While this aftereffect is typically measured in unisensory trials, it could also shape responses during subsequent multisensory trials and thereby contribute to the numerically observed dependency of the ventriloquism bias on preceding discrepancies. That is, the aftereffect induced by (and measured in) one AV-A pair could shape the ventriloquism bias in a following AV trial. To probe whether this is the case, we implemented a more extended model for the ventriloquism effect that, in addition to the spatial discrepancies, also included the aftereffects in preceding trial-pairs. Specifically, we modelled the ventriloquism bias in trial-pair d based on the discrepancies in pairs b,c, and d and on the aftereffect in pairs a,b and c. We then used variance partitioning to probe whether the aftereffect contributes uniquely to shaping subsequent ventriloquism biases and whether it contributes common variance that overlaps with that explained by the spatial discrepancies. Overlapping variance could arise if an apparent history dependence of the bias is shaped by the preceding aftereffects. Unique variance would point to an influence of the aftereffect that is separate from the multisensory discrepancy, for example if the aftereffect constitutes a genuine modality-specific bias.

The results suggest that the aftereffect did not contribute significant unique or shared variance to the ventriloquism effect (Figure 4). As expected from the above, the immediate spatial discrepancy (termed ‘Δva’) explained a considerable variance (0.324 [0.280 0.370]), while the contribution of the preceding discrepancies was negligible (‘ΔvaPast’: −0.002 [−0.003 −0.000]). Importantly, the unique contribution of the aftereffect was only marginal (‘ae’: 0.005 [0.003 0.008]) as was its common contribution with the immediate or preceding discrepancies (0.001 [0 0.002], and 0.003 [0.002 0.005]). Hence, any residual dependency of the ventriloquism bias that is reflected in these datasets is only marginally explained by specific contributions of previous spatial arrangements or the aftereffect.

**Figure 4:**
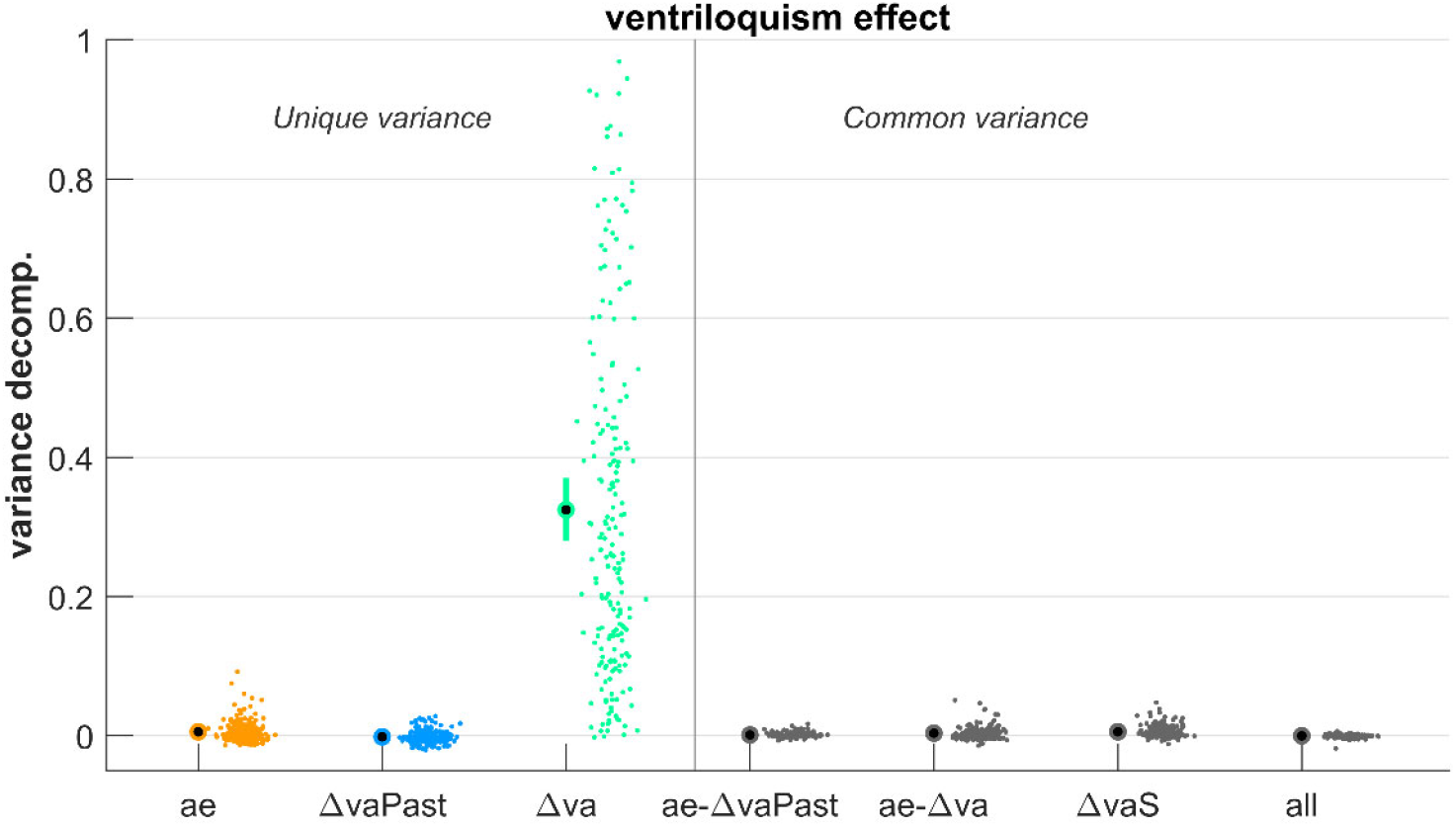
Modelling the ventriloquism bias against spatial discrepancies and previous aftereffects. We modelled the ventriloquism bias against the spatial discrepancy in the immediate AV trial-pair (labelled Δva) and two preceding trial pairs (labelled ΔvaPast). In addition, we included the aftereffect in three preceding trial pairs (labelled ae). These models featured both linear and non-linear dependencies on discrepancy and a linear dependency on the aftereffect. The graph shows the results from a variance partitioning analysis, with unique variances in colour and common variances in grey (here ae-ΔvaPast/ae-Δva are the variances shared by the aftereffect and the preceding / immediate discrepancies, ΔvaS denotes the variance shared by immediate and preceding discrepancies and all refers to the variance common to all predictor groups). Common variances were all negligible with mean values close to zero and/or 99% CIs reaching zero. Dots reflect individual datasets (i.e. participants, n=205), circles the overall mean and error bars the 99% bootstrap confidence interval.

## Discussion

Multisensory integration and recalibration are two phenomena by which perception deals with discrepant signals from our environment. Such discrepancies can be dynamic and may feature predictable and unpredictable components. Our results show that the spatial ventriloquism bias - reflecting integration - is shaped almost exclusively by the immediate multisensory discrepancy and not by those presented in preceding trials. In contrast, the ventriloquism aftereffect - reflecting recalibration - scales in a cumulative manner with these discrepancies. Hence the history of multisensory experience allows dissociating these two biases, supporting the conclusion that the ventriloquism aftereffect is not an obligatory and direct consequence of the integration of discrepant multisensory signals.

### Multisensory integration as adaptive process

Our data support the notion that multisensory integration reduces discrepant sensory estimates between seemingly redundant signals (Shams and Beierholm, 2010; Odegaard et al., 2015; Rohe and Noppeney, 2015b). In the large data sample analysed here, the ventriloquism bias was driven mostly by the audio-visual discrepancy experienced in the current trial, while those experienced in the preceding multisensory trials had only a minimal influence, at best. Still, this result does not rule out that the overall integration process also reflects the history of the sensory environment. Integration scales with the relative unisensory reliabilities (Ernst and Bulthoff, 2004; Angelaki et al., 2009) and our brain keeps track of these in a leaky manner (Beierholm et al., 2020). This renders the integration weights assigned to the individual signals history dependent. Similarly, our brain tracks the presumed causal relations between multisensory signals, and the momentary estimate of whether two signals are deemed causally related depends on the recent exposure to discrepant stimuli (Odegaard et al., 2017; Rohe et al., 2019). This history dependence of a common-cause prior provides a route for the past multisensory experience to enter the general integration process, similar to generic Bayesian models of perception in which the previous trial’s posterior influences the next trials prior (van Bergen and Jehee, 2019).

However, our data show that previous multisensory discrepancies do not directly shape the integration bias. That is, the sign or magnitude of previously experienced discrepancies have very limited direct predictive power for the subsequent integration bias. This conclusion is in line with a previous study that was based on a much smaller sample and relied on an experimental design featuring only one magnitude of spatial discrepancy (Bosen et al., 2017). In that study, the audio-visual stimuli were either discrepant or co-localized, which may influence the manner in which multisensory causal relations are estimated. In contrast, in the present experiments both the degree and direction of spatial discrepancies were unpredictable, demonstrating the lack of history-dependence of the integration bias in a more variable and possibly more natural context.

One could argue that any influence of previous trials on integration emerges in the form of spatial recalibration, whereby the previous trial-pair shifts auditory representations that bias the integration in subsequent AV trials. To test for such a propagation of auditory recalibration across trial-pairs, we probed the predictive power of the aftereffect bias on the subsequent ventriloquism bias. However, this contribution was numerically marginal compared to the multisensory discrepancy in the AV trial of interest, suggesting that contributions of cumulative recalibration to the ventriloquism bias are also negligible.

Across the three types of data analyses presented here, the numeric evidence for a history dependence of integration differed. The mixed effect model across all datasets was clearly in favour of no history-dependence and the participant-wise models attributed negligible unique variance to previous trials. This suggests that the slight numeric differences in the ventriloquism effect between sequences of trial-pairs with same or changing discrepancies in Figure 2 possibly result from comparing the specific selected trials and does not generalize to the full datasets as analysed using linear models.

### Multisensory recalibration accumulates over time

Our results show a clear cumulative effect of preceding multisensory discrepancies on the aftereffect. This observation is in line with the known adaptive nature of the aftereffect, which is classically probed following several minutes of exposure to discrepant audio-visual stimuli (Radeau and Bertelson, 1974; Recanzone, 1998; Bruns and Roder, 2015; Bosen et al., 2018; Bruns, 2019; Bruns and Röder, 2019). However, recent studies have shown that this aftereffect emerges also on a trial-wise basis (Wozny and Shams, 2011b; Bosen et al., 2017; Park and Kayser, 2019) and some have shown that the aftereffect accumulates over a series of up to 5 (Wozny and Shams, 2011b; Mendonça et al., 2015) or twenty trials (Frissen et al., 2012; Bosen et al., 2017).

Our results extend those studies in a number of ways. First, most studies probing the influence of immediately preceding trials relied on a fixed magnitude of the spatial discrepancy, or even a fixed sign (Mendonça et al., 2015; Bosen et al., 2017). Such a high predictability may foster the cumulative nature of the aftereffect and may engage additional learning mechanisms not present in more naturalistic environments involving pseudo-random and unpredictable discrepancies as used here. Second, the locations of the auditory stimuli in AV and subsequent A trials were either the same or limited to within a 10°range in previous work (Mendonça et al., 2015; Bosen et al., 2017). In contrast, in the present experiments they varied to a much larger extend, requiring recalibration to generalize over a larger proportion of the auditory space. Hence, the simple repetition of previous judgements that participants may have used in previous studies was not possible in this design. Third, because of their design, these studies did not directly quantify the independent contribution of the preceding trials, which the variance decomposition here does. Furthermore, eye movements or the direction of fixations may confound trial-wise recalibration (Razavi et al., 2007; Dobreva et al., 2012; Park and Kayser, 2019), which emerges in a mixture of head- and eye-centered reference frames (Kopco et al., 2009). To avoid such confounding effects, the present study relied on very short auditory and visual signals, making it unlikely that participants moved their eyes during stimulus presentation (see also discussions in (Park and Kayser, 2019, 2021) and (Bosen et al., 2017)).

Similar to most previous studies the present experiments relied on a predictable trial structure in which AV trials were always followed by A trials. This repetitive nature may strengthen the influence of preceding multisensory discrepancies, whose occurrence in the trial sequence can to some degree be predicted. However, this feature is unlikely to solely explain the history dependence of the aftereffect, given that a similar cumulative property is also observed with pseudo-randomized sequences of trials (Wozny and Shams, 2011b).

### How are integration and recalibration linked?

One view stipulates that recalibration is driven by the discrepancy between integrated multisensory signals and a subsequent unisensory stimulus, rendering recalibration directly dependent on the outcome of integration (Zaidel et al., 2013; Bruns, 2019; Noppeney, 2021). If two multisensory signals are deemed sufficiently discrepant to unlikely originate from a common source, the outcome of multisensory causal inference should emphasize the task-relevant unisensory estimate, leaving no multisensory bias to drive recalibration. Hence, in this view integration becomes a prerequisite for recalibration, similar to development where integration seems to develop prior to recalibration (Rohlf et al., 2020). While our data cannot rule out that integration per se is required for recalibration to emerge, our data speak against the hypothesis that both processes are directly linked by a similar dependency on preceding multisensory discrepancies.

An alternative view holds that recalibration is shaped by the believe in a modality-specific bias (Di Luca et al., 2009; Block and Bastian, 2011; Burr et al., 2011; Zaidel et al., 2013; Noppeney, 2021). This believe may be shaped by multiple factors, including judgements about the causal relation of sensory signals as one of many factors. As a result, both integration and recalibration tend to correlate across experimental manipulations and the immediate multisensory discrepancy. However, a residual ventriloquism aftereffect emerges also when auditory and visual signals are not judged as originating from a common source (Wozny and Shams, 2011b), when obviously not originating from a common location (Radeau and Bertelson, 1974) and when attention is directed towards task-unrelated visual stimuli (Eramudugolla et al., 2011). Hence, recalibration is not directly contingent on multisensory signals to be judged as relating to the same object. Our results support this view and speak in favour of distinct functional roles of integration and recalibration that are shaped by the immediate multisensory discrepancy as just one of many factors.

The collective evidence is in line with a model of Bayesian causal inference that shapes multisensory perception in general, but which affects integration and recalibration via distinct mechanisms (Noppeney, 2021). Unpredictable discrepancies between auditory and visual stimuli are reduced by multisensory integration, and the brain continuously updates the a priori believe in a common cause of multisensory signals based on the recently experienced discrepancies (Kording et al., 2007; Beierholm et al., 2020; Shams and Beierholm, 2022). At the same time, the representation of auditory space is continuously updated in a leaky manner to compensate for short-term changes in audio-visual disparities and to minimize apparent localization errors (Kopco et al., 2009; Wozny and Shams, 2011a). The respectively underlying estimates of audio-visual disparities may be indirectly shaped by the common cause prior, but it is possible that distinct time scales of this are relevant for integration and recalibration. In particular, for integration that primarily depends on the instantaneous bimodal signals, the common cause evidence should be based on a short time scale, whereas for recalibration that depends on a series of bimodal signals it should be based on a longer time scale. Testing this prediction may not be easily feasible with the very brief stimuli used here, but for example paradigms involving audio-visual motion or visuo-motor paradigms involving hand movements and manipulated visual feedback could allow this (Debats et al., 2017; Debats and Heuer, 2018).

## Acknowledgements

This study was funded by the Deutsche Forschungsgemeinschaft (DFG KA2661/2-1). The authors declare that they have no conflict of interests. We would like to thank several undergraduate students and research assistants for their help during data collection.

## Data and code

The full data analysed here and the respective code are available here: https://github.com/Hame-p/recal_history

## References

Angelaki DE, Gu Y, DeAngelis GC (2009) Multisensory integration: psychophysics, neurophysiology, and computation. Curr Opin Neurobiol 19:452–458.

Badde S, Navarro KT, Landy MS (2020) Modality-specific attention attenuates visual-tactile integration and recalibration effects by reducing prior expectations of a common source for vision and touch. Cognition 197:104170.

Beierholm U, Rohe T, Ferrari A, Stegle O, Noppeney U (2020) Using the past to estimate sensory uncertainty. eLife 9.

Benjamini Y, Yu B (2013) THE SHUFFLE ESTIMATOR FOR EXPLAINABLE VARIANCE IN FMRI EXPERIMENTS. The Annals of Applied Statistics 7:2007–2033.

Block HJ, Bastian AJ (2011) Sensory weighting and realignment: independent compensatory processes. J Neurophysiol 106:59–70.

Bosen AK, Fleming JT, Allen PD, O’Neill WE, Paige GD (2017) Accumulation and decay of visual capture and the ventriloquism aftereffect caused by brief audio-visual disparities. Exp Brain Res 235:585–595.

Bosen AK, Fleming JT, Allen PD, O’Neill WE, Paige GD, Bosen AK, Chatterjee M (2018) Multiple time scales of the ventriloquism aftereffect. PLoS One 13:e0200930.

Bruns P (2019) The Ventriloquist Illusion as a Tool to Study Multisensory Processing: An Update. Front Integr Neurosci 13:51.

Bruns P, Roder B (2015) Sensory recalibration integrates information from the immediate and the cumulative past. Sci Rep 5:12739.

Bruns P, Röder B (2019) Repeated but not incremental training enhances cross-modal recalibration. J Exp Psychol Hum Percept Perform 45:435–440.

Burr D, Binda P, Gori M (2011) 173Multisensory Integration and Calibration in Adults and in Children. In: Sensory Cue Integration (Trommershäuser J, Kording K, Landy MS, eds), p 0: Oxford University Press.

Debats NB, Heuer H (2018) Optimal integration of actions and their visual effects is based on both online and prior causality evidence. Sci Rep 8:9796.

Debats NB, Ernst MO, Heuer H (2017) Perceptual attraction in tool use: evidence for a reliability-based weighting mechanism. J Neurophysiol 117:1569–1580.

Di Luca M, Machulla TK, Ernst MO (2009) Recalibration of multisensory simultaneity: cross-modal transfer coincides with a change in perceptual latency. J Vis 9:7.1–16.

Dobreva MS, O’Neill WE, Paige GD (2012) Influence of age, spatial memory, and ocular fixation on localization of auditory, visual, and bimodal targets by human subjects. Exp Brain Res 223:441–455.

Eramudugolla R, Kamke MR, Soto-Faraco S, Mattingley JB (2011) Perceptual load influences auditory space perception in the ventriloquist aftereffect. Cognition 118:62–74.

Ernst MO, Bulthoff HH (2004) Merging the senses into a robust percept. Trends Cogn Sci 8:162–169.

Fethney J (2010) Statistical and clinical significance, and how to use confidence intervals to help interpret both. Australian critical care: official journal of the Confederation of Australian Critical Care Nurses 23:93–97.

Freckleton RP (2002) On the misuse of residuals in ecology: regression of residuals vs. multiple regression. Journal of Animal Ecology 71:542–545.

Frissen I, Vroomen J, de Gelder B (2012) The aftereffects of ventriloquism: the time course of the visual recalibration of auditory localization. Seeing Perceiving 25:1–14.

Gelman A, Hwang J, Vehtari A (2014) Understanding predictive information criteria for Bayesian models. Statistics and Computing 24:997–1016.

Greenland S, Senn SJ, Rothman KJ, Carlin JB, Poole C, Goodman SN, Altman DG (2016) Statistical tests, P values, confidence intervals, and power: a guide to misinterpretations. European journal of epidemiology 31:337–350.

Kopco N, Lin IF, Shinn-Cunningham BG, Groh JM (2009) Reference frame of the ventriloquism aftereffect. J Neurosci 29:13809–13814.

Kording KP (2014) Bayesian statistics: relevant for the brain? Current Opinion in Neurobiology 25:130–133.

Kording KP, Beierholm U, Ma WJ, Quartz S, Tenenbaum JB, Shams L (2007) Causal inference in multisensory perception. PLoS ONE 2:e943.

Mendonça C, Escher A, van de Par S, Colonius H (2015) Predicting auditory space calibration from recent multisensory experience. Exp Brain Res 233:1983–1991.

Nimon K, Lewis M, Kane R, Haynes RM (2008) An R package to compute commonality coefficients in the multiple regression case: an introduction to the package and a practical example. Behav Res Methods 40:457–466.

Noppeney U (2021) Perceptual Inference, Learning, and Attention in a Multisensory World. Annu Rev Neurosci 44:449–473.

Odegaard B, Wozny DR, Shams L (2015) Biases in Visual, Auditory, and Audiovisual Perception of Space. PLoS Comput Biol 11:e1004649.

Odegaard B, Wozny DR, Shams L, Odegaard B, Shams L (2017) A simple and efficient method to enhance audiovisual binding tendencies The Brain’s Tendency to Bind Audiovisual Signals Is Stable but Not General. PeerJ 5:e3143.

Park H, Kayser C (2019) Shared neural underpinnings of multisensory integration and trial-by-trial perceptual recalibration in humans. eLife 8.

Park H, Kayser C (2020) Robust spatial ventriloquism effect and trial-by-trial aftereffect under memory interference. Sci Rep 10:20826.

Park H, Kayser C (2021) The Neurophysiological Basis of the Trial-Wise and Cumulative Ventriloquism Aftereffects. J Neurosci 41:1068–1079.

Park H, Kayser C (2022) The context of experienced sensory discrepancies shapes multisensory integration and recalibration differently. Cognition 225:105092.

Park H, Nannt J, Kayser C (2021) Sensory-and memory-related drivers for altered ventriloquism effects and aftereffects in older adults. Cortex 135:298–310.

Radeau M, Bertelson P (1974) The after-effects of ventriloquism. The Quarterly journal of experimental psychology 26:63–71.

Raftery AE (1995) Bayesian Model Selection in Social Research. Sociological Methodology 25:111–163.

Razavi B, O’Neill WE, Paige GD (2007) Auditory spatial perception dynamically realigns with changing eye position. J Neurosci 27:10249–10258.

Recanzone GH (1998) Rapidly induced auditory plasticity: the ventriloquism aftereffect. Proc Natl Acad Sci U S A 95:869–875.

Recanzone GH (2009) Interactions of auditory and visual stimuli in space and time. Hear Res 258:89–99.

Rohe T, Noppeney U (2015a) Cortical Hierarchies Perform Bayesian Causal Inference in Multisensory Perception. PLoS Biol 13: e1002073

Rohe T, Noppeney U (2015b) Sensory reliability shapes perceptual inference via two mechanisms. J Vis 15:22.

Rohe T, Ehlis AC, Noppeney U (2019) The neural dynamics of hierarchical Bayesian causal inference in multisensory perception. Nature communications 10:1907.

Rohlf S, Bruns P, Röder B (2021) The Effects of Cue Reliability on Crossmodal Recalibration in Adults and Children. Multisens Res:1–19.

Rohlf S, Li L, Bruns P, Röder B (2020) Multisensory Integration Develops Prior to Crossmodal Recalibration. Curr Biol 30:1726–1732.e1727.

Rouder JN, Speckman PL, Sun D, Morey RD, Iverson G (2009) Bayesian t tests for accepting and rejecting the null hypothesis. Psychon Bull Rev 16:225–237.

Seibold DR, McPhee RD (1979) COMMONALITY ANALYSIS: A METHOD FOR DECOMPOSING EXPLAINED VARIANCE IN MULTIPLE REGRESSION ANALYSES. Human Communication Research 5:355–365.

Shams L, Beierholm UR (2010) Causal inference in perception. Trends Cogn Sci 14:425–432.

Shams L, Beierholm U (2022) Bayesian causal inference: A unifying neuroscience theory. Neurosci Biobehav Rev 137:104619.

van Bergen RS, Jehee JFM (2019) Probabilistic Representation in Human Visual Cortex Reflects Uncertainty in Serial Decisions. J Neurosci 39:8164–8176.

Wagenmakers EJ (2007) A practical solution to the pervasive problems of p values. Psychon Bull Rev 14:779–804.

Wozny DR, Shams L (2011a) Computational characterization of visually induced auditory spatial adaptation. Front Integr Neurosci 5:75.

Wozny DR, Shams L (2011b) Recalibration of auditory space following milliseconds of cross-modal discrepancy. J Neurosci 31:4607–4612.

Zaidel A, Ma WJ, Angelaki DE (2013) Supervised Calibration Relies on the Multisensory Percept. Neuron.

